# CHAMP1 Complex Promotes Heterochromatin Assembly and Reduces Replication Stress

**DOI:** 10.1101/2025.08.05.667964

**Authors:** Feng Li, Amira Elbakry, Felix Y. Zhou, Tianpeng Zhang, Ramya Ravindranathan, Huy Nguyen, Aleem Syed, Lifang Sun, Sirisha Mukkavalli, Roger A. Greenberg, Alan D. D’Andrea

## Abstract

Replication stress is a major driver of genomic instability and a hallmark of cancer cells. Although dynamic heterochromatin remodeling has been implicated in replication stress response, the precise mechanisms remain unclear. Here, we identify the CHAMP1 complex, composed of CHAMP1, POGZ, HP1α, and the H3K9 methyltransferase SETDB1, as a critical regulator of heterochromatin assembly at stalled replication forks. Upon replication stress, the CHAMP1 complex is recruited to stalled forks where it facilitates H3K9me3 deposition, creating a repressive chromatin environment that shields replication forks from MRE11-mediated degradation. The complex promotes the recruitment of the origin recognition complex (ORC) to sites of replication stress, such as the telomeric heterochromatin in alternative lengthening of telomeres (ALT)-positive tumor cells, thereby supporting efficient telomeric DNA replication. Loss of CHAMP1 disrupts ORC2 recruitment and impairs fork restart, leading to increased micronuclei formation and heightened sensitivity to replication stress. Notably, CHAMP1 deficiency induces synthetic lethality with FANCM inhibition in ALT-positive tumor cells, and the CHAMP1 complex is essential for the survival of CCNE1-amplified ovarian cancers. These findings uncover a chromatin-based mechanism of replication fork stabilization and suggest that CHAMP1 may represent a candidate therapeutic vulnerability in cancers with elevated replication stress.

## INTRODUCTION

Genomic instability is a hallmark of cancer and is often driven by replication stress (RS)— a condition in which replication forks stall or collapse due to obstacles during DNA synthesis. RS is often triggered by a variety of oncogenic events, including overexpression of MYC or Cyclin E, loss of the p53-mediated G1/S checkpoint, or deficiencies in DNA repair pathways ^1–4^. These replication stress events result in the cellular accumulation of free single-strand DNA, triggering activation of the ATR/CHK1 pathway, which slows cell cycle progression and alleviates replication stress ^5^.

Replication stress can also be localized to specific regions of the genome and is amplified in specific cellular contexts. For instance, in about 10-15% human cancers, telomere length is maintained by the alternative lengthening of telomere (ALT) mechanism^6^ ^7^ which bypasses the need for telomerase-mediated telomere lengthening. ALT-positive tumor cells are characterized by high local replication stress at telomeres and rely heavily on homology-directed repair (HDR) and DNA damage response pathways, including the ATR/CHK1 pathway ^8^ and the Fanconi anemia pathway ^9^. Notably, ALT tumor cells are highly sensitive to inhibitors of the FANCM translocase enzyme ^10^ ^11^ ^12^.

Recent studies have implicated the CHAMP1 protein complex in regulating chromatin assembly at ALT telomeres ^13^. This complex recruits the methyltransferase, SETDB1, to sites of heterochromatin assembly, such as telomeres and centromeres and increases the local level of H3K9me3 at these sites ^13^. This process is required for the clustering of telomeres, known to promote homology-directed DNA repair^14^ and is critical for the recruitment of homologous recombination proteins to these sites and the local repair of DNA double-strand breaks ^13^. The CHAMP1 complex also promotes homology-directed repair at other DSBs in euchromatin, by binding to REV7 and reducing the level of the SHLD protein complex ^15^ ^16^. The SHLD complex normally blocks DSB end resection and thereby suppresses homologous recombination ^17–21^.

Beyond their roles in chromatin regulation and DNA repair, CHAMP1 and POGZ are clinically relevant genes. Patients with *de novo* heterozygous mutations in the *CHAMP1 gene*, known as CHAMP1 syndrome, present with a neurodevelopmental disorder characterized by intellectual disability, behavioral symptoms, and distinct dysmorphic features ^22–26^. A recent study indicates that patients with CHAMP1 syndrome develop acute myeloid leukemia, suggesting a broader role of the CHAMP1 complex in reducing replication stress and maintaining the genomic stability of hematopoietic stem and progenitor cells ^22^. Interestingly*, de novo* heterozygous mutations in the *POGZ* gene also result in a rare but highly related neurodevelopmental syndrome called the White Sutton Syndrome (WSS) ^27–35^. These human syndromes underscore the physiological importance of CHAMP1/POGZ in regulating chromatin and maintaining genome integrity. How this complex protects against replication stress in proliferating cells remains poorly understood.

The Origin Recognition Complex (ORC) is a heteromeric six-subunit protein complex that plays a central role in eukaryotic DNA replication by binding to replication origin sites and serving as a scaffold for the assembly of other key initiation factors (reviewed in ^36^). Recent studies have identified a role of ORC6 in reducing replication stress by promoting efficient and accurate DNA replication ^37,38^. However, it remains unclear whether other components of the ORC complex contribute to this function. ORC also plays a critical role at telomeres, where it initiates DNA replication at chromosome ends to ensure proper telomere duplication and prevent telomere shortening ^39^ ^40^. ORC is recruited to telomeres by the shelterin protein TRF2 ^41^ ^42^ and by HP1 ^43^. However, whether ORC is recruited to ALT telomeres in response to high replication stress remains unclear.

Here, we show that the CHAMP1 complex plays a critical role in reducing replication stress in human cancer cells. Loss of the CHAMP1 complex sensitizes cells to replication stress-inducing agents such as hydroxyurea (HU) and induces compensatory mechanisms, including the ATR/CHK1 pathway and the FA pathway. Loss of the CHAMP1 complex results in recruitment to telomeres of proteins involved in the FA pathway, including the ATPase, FANCM to telomeres. Strikingly, we discovered that CHAMP1 recruits ORC2 to telomeres and stalled forks. Loss of ORC2 resulted in increased sensitivity to HU. Together, our findings reveal a novel function of the CHAMP1 complex in orchestrating H3K9me3-mediated ORC2 recruitment during replication stress and demonstrate that the CHAMP1 complex is essential in cancer cells with high levels of replication stress.

## RESULTS

### CHAMP1 complex relieves replication stress and prevents micronuclei formation

To determine the role of the CHAMP1 complex in alleviating replication stress, we conducted a series of assays in CHAMP1 knockout (KO) RPE1^p53-/-^ cells (**Figure 1**). CHAMP1-deficient cells exhibited increased sensitivity to replication stress-inducing agents, including hydroxyurea (HU) (**Figure 1A**), aphidicolin (**Figure 1B**), and cisplatin (**Figure 1C**). Similar sensitivity was observed in POGZ-deficient RPE cells, consistent with POGZ’s role as a CHAMP1 interactor (**Supplemental Figure 1**). Upon HU treatment, CHAMP1 KO cells exhibited increased levels of replication stress markers, including phosphorylated KAP1 and CHK1, in a time-dependent manner (**Figure 1D**), indicating an exacerbated replication stress response. These cells were also more sensitive to ATR inhibition (**Figure 1E**), which impairs the replication stress response ^4,44^. Furthermore, CHAMP1 depletion significantly increased chromosomal bridges and micronuclei formation (**Figure 1F, G**), indicative of replication catastrophe and genome instability ^4,45,46^.

**Fig. 1.**
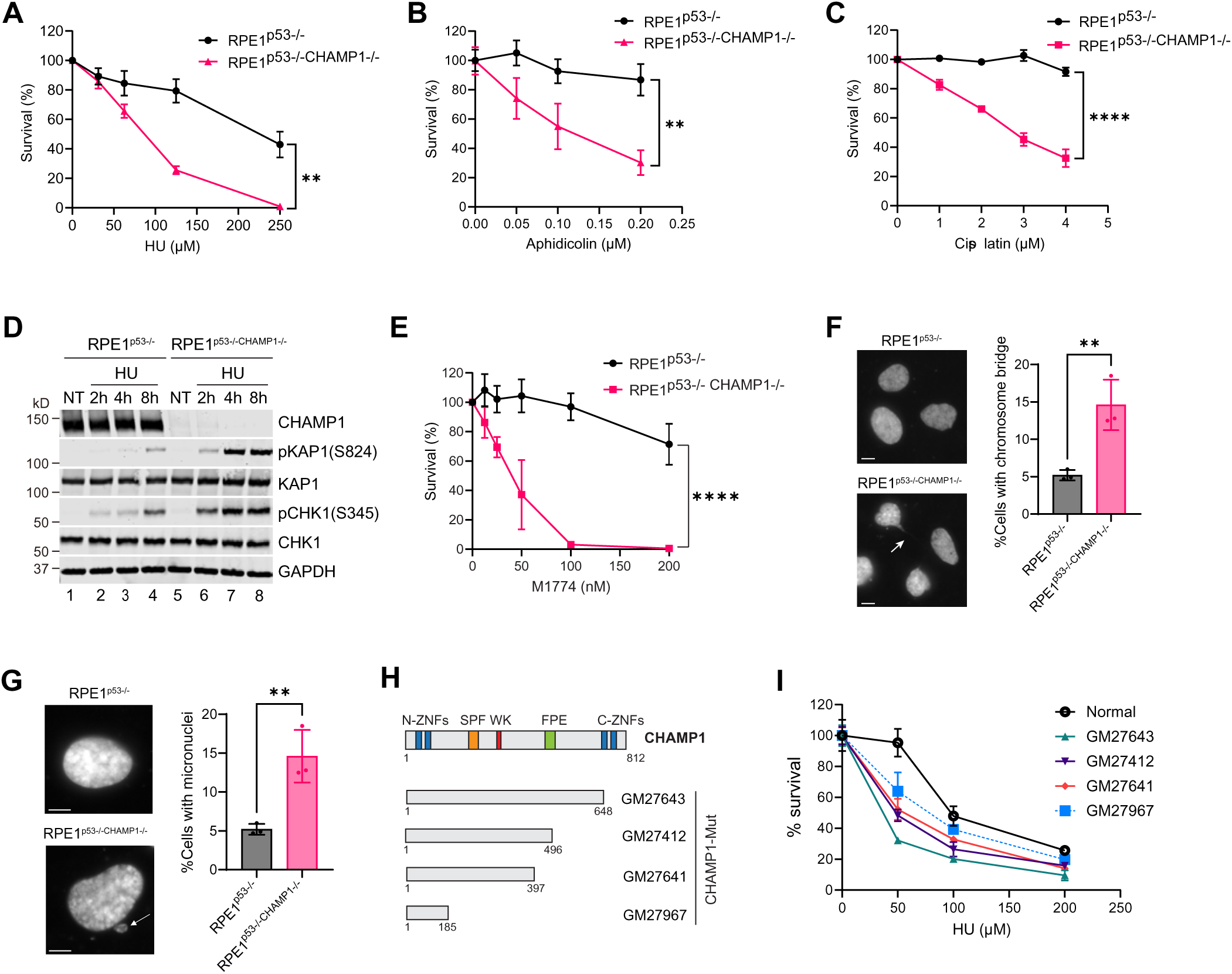
Loss of CHAMP1 results in heightened sensitivity to replication stress and increased micronuclei formation. **A-C.** Colony survival plots of RPE1-p53-/- and RPE1-p53-/- CHAMP1-/- cells treated with different concentrations of hydroxyurea (HU) (**A**), aphidicolin (**B**), or cisplatin (**C**) for 10 days. **D.** Western blot analysis of replication stress response signaling from RPE1-p53-/- and RPE1-p53-/- CHAMP1-/- cells after 2, 4, or 8 hours 4mM HU treatment. The Western blots are representative of two independent experiments. **E.** Colony survival plots of RPE1-p53-/- and RPE1-p53-/- CHAMP1-/- cells treated with different concentrations of ATR inhibitor M1774 for 10 days. **F**. (left) Representative images of cells with and without a chromosome bridge. Arrow points to a chromosome bridge. DAPI was used to stain the nuclei. Scale bar, 5 µm. (right) Percentage of interphase cells containing a chromosome bridge is presented as mean values ±SD, n=3 independent experiments. More than 100 cells were counted for each experiment. P values were calculated using a two-tailed Student’s t test. **G**. (left) Representative images of cells with and without micronuclei. Arrow points to a micronuclei. DAPI was used to stain the nuclei. Scale bar, 5 µm. (right) Percentage of interphase cells containing micronuclei are presented as mean values ±SD, n=3 independent experiments. More than 100 cells were counted for each experiment. P values were calculated using a two-tailed Student’s t test. **H.** Schematic of CHAMP1 mutants from EBV-immortalized lymphoblast cell lines derived from several CHAMP1 patients (Coriell). **I.** 3-day cytotoxicity analysis of the normal and CHAMP1 patient lymphoblast cells treated with various doses of HU. Cell viability was detected by CellTiter Glo (Promega). Error bars indicate SD, n=3 independent experiments.

Importantly, EBV-immortalized lymphoblastoid cell lines (Coriell) derived from CHAMP1 syndrome patients—who carry a heterozygous CHAMP1 mutation—express truncated CHAMP1 proteins lacking the C-terminal HP1α-binding domain ^13^ (**Figure 1H**). These mutant proteins may act dominantly negative or destabilize the protein, resulting in haploinsufficiency and disrupt heterochromatin assembly in these heterozygous cells. Notably, these patient-derived cells are also hypersensitive to HU (**Figure 1I**), suggesting a compromised replication stress response. Collectively, these findings demonstrate that the CHAMP1 complex is essential for maintaining genome integrity under replication stress conditions.

### CHAMP1 complex promotes replication fork stability and facilitates restart under stress

To further investigate the role of the CHAMP1 complex in the cellular response to replication stress, we first examined its localization to replication forks. Using the SIRF (in situ analysis of protein interactions at DNA replication forks) assay—which combines proximity ligation chemistry with EdU (5-Ethynyl-2’-deoxyuridine) labeling of nascent DNA—we confirmed that CHAMP1 is recruited to stalled replication forks by hydroxyurea (HU) treatment (**Figure 2A, B**). Complementary experiments using native IdU (5-iodo-2′-deoxyuridine) staining revealed that HU treatment elevates single-stranded DNA (ssDNA) levels in RPE1^p53-/-^ cells, which is a marker of replication stress ^47^. Notably, CHAMP1-deficient (CHAMP1-/-) cells exhibited higher baseline ssDNA, which was further amplified by HU treatment (**Figure 2C, D**), indicating an inherent vulnerability to replication stress in the absence of CHAMP1.

**Fig. 2.**
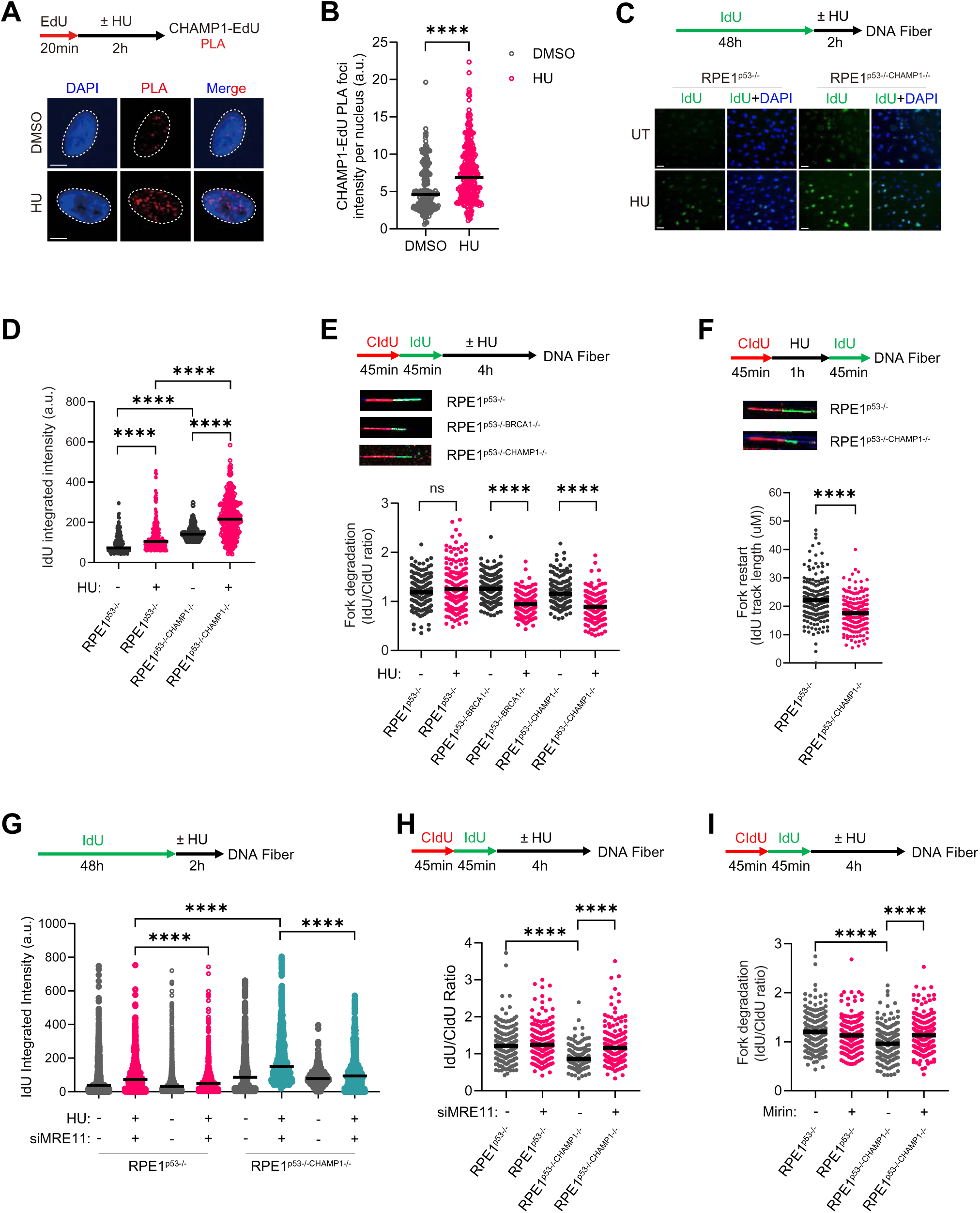
Loss of CHAMP1 results in replication fork instability and restart defect. **A.** Representative images of proximity ligation assay (PLA) showing CHAMP1 localization at replication sites (CHAMP1-EdU PLA, red). Nuclei were stained with DAPI (blue). Scale bar, 5 µm. Cells were pulse-labeled with EdU (20 μM) for 20 min, followed by treatment with either 4 mM hydroxyurea (HU) or DMSO for 2 hours. **B.** Distribution of the total intensity of all CHAMP1-EdU PLA spots per nucleus in RPE1^p53-/-^ cells treated with HU or DMSO. More than 50 cells were counted for each of the three independent experiments. P values were calculated using the Mann-Whitney test. **C.** Left: Representative images of IdU staining without denaturation, indicating regions of single-stranded DNA (ssDNA). Nuclei were counterstained with DAPI (blue), Scale bar, 20 µm. Cells were incubated with IdU for 48 hours and subsequently treated with either 4 mM HU or DMSO for 2 hours. **D.** Distribution of the total intensity of IdU per nucleus in RPE1^p53-/-^ or RPE1^p53-/-CHAMP1-/-^ cells treated with HU or DMSO. More than 50 cells were counted for each of the three independent experiments. P values were calculated using Mann-Whitney test. **E.** Top: Representative images of DNA fibers illustrating replication fork degradation. Cells were sequentially labeled with CIdU and IdU for 45 minutes each, followed by treatment with either 4 mM HU or DMSO for 4 hours before performing the DNA fiber assay. Bottom: Quantification of replication fork degradation, shown as the ratio of IdU to CIdU tract lengths. Each dot represents an individual fiber, with at least 200 fibers analyzed per condition. Statistical significance was determined using the Mann–Whitney U test. **F.** Top: Representative DNA fiber images showing replication fork restart. Cells were labeled with CIdU, treated with 4 mM HU for 1 hour, then released into IdU for 45 minutes before performing the DNA fiber assay. Bottom: Quantification of replication fork restart, shown as the IdU tract lengths. Each dot represents an individual fiber, with at least 200 fibers analyzed per condition. Horizontal bars indicate medians. Statistical significance was determined using the Mann–Whitney U test. **G.** RPE1^p53-/-^ or RPE1^p53-/-CHAMP1-/-^ cells were treated with siControl or siMRE1, and then incubated with IdU for 48 hours, followed by treatment with either 4 mM HU or DMSO for 2 hours. The total intensity of IdU per nucleus was quantified. For each of three independent experiments, more than 50 cells were analyzed. P values were calculated using Mann-Whitney test. **H-I.** RPE1^p53-/-^ or RPE1^p53-/-CHAMP1-/-^ cells treated with siMRE11 (**H**) or the MRE11 inhibitor Mirin (**I**), then sequentially labeled with CIdU and IdU for 45 minutes each. After labeling, cells were exposed to 4 mM HU or DMSO for 4 hours before DNA fiber assays were performed. Replication fork degradation was quantified by calculating the IdU/CIdU tract length ratio. At least 200 fibers were analyzed per condition. Statistical significance was determined using the Mann–Whitney U test.

Further analysis using DNA fiber assays provided insights into replication fork dynamics. Upon HU treatment, CHAMP1-/- cells revealed a marked decrease in the IdU/CldU tract length ratio (**Figure 2E**), indicating increased nascent fork degradation—an observation that mirrors the established role of BRCA1 in fork protection ^48^. Similar fork degradation was observed in POGZ-deficient RPE-1 cells (**Supplemental Figure 2A**), and POGZ knockdown in CHAMP1 knockout cells did not further exacerbate the phenotype (**Supplemental Figure 2A**), suggesting that CHAMP1 and POGZ function together as a complex to regulate stalled replication fork stability. Moreover, CHAMP1 loss impaired replication fork restart, as shown by decreased frequency and shorter length of restarted forks in a modified DNA fiber assay (**Figure 2F**). These results suggest that CHAMP1 deficiency not only enhances fork degradation but also impairs the efficient restart of stalled replication forks.

To assess whether MRE11 mediates fork degradation in the absence of CHAMP1, we first assessed its recruitment to stalled replication forks. As expected, MRE11 was enriched at HU-treated stalled replication forks (**Supplemental Figure 2B, C**). Notably, CHAMP1-depleted cells showed a marked increase in MRE11 accumulation at stalled replication forks (**Supplemental Figure 2B, C**), suggesting enhanced nuclease activity in the absence of CHAMP1. Depletion of MRE11 using siRNA rescued the ssDNA accumulation and fork degradation phenotype in CHAMP1 knockout cells (**Figure 2G, H**). Similarly, treatment with Mirin, a selective MRE11 inhibitor, reversed the degradation (**Figure 2I**). These findings indicate that MRE11 is the primary nuclease responsible for fork degradation in CHAMP1-deficient cells. Taken together, the CHAMP1 complex appears to play a critical role in protecting replication forks from MRE11-dependent processing under replication stress.

### CHAMP1 complex induces H3K9me3 accumulation at stalled replication forks

We previously demonstrated that CHAMP1 and POGZ form a core complex that assembles and maintains heterochromatin via interaction with HP1α and the methyltransferase, SETDB1 ^13^. The complex localizes to H3K9me3-marked regions ^13^. *De novo* H3K9me3 formation at stalled replication forks is known to be crucial for replication fork integrity ^49^. To investigate whether the CHAMP1 complex is involved in this process, we first examined whether replication stress enhances CHAMP1 complex assembly and promotes H3K9me3 accumulation. Interestingly, brief treatment with HU or aphidicolin increased the interaction between CHAMP1 and its associated partners POGZ, HP1α, and H3K9me3 (**Figure 3A**), suggesting that replication stress induces CHAMP1 complex formation. We next employed the SIRF assay to detect H3K9me3 at nascent DNA after HU treatment. Consistent with previous studies ^14^, HU treatment increased H3K9me3 and H3K9me2 at the stressed replication sites (**Figure 3B, C**). Notably, both histone marks were significantly reduced in CHAMP1 and POGZ knockout cells (**Figure 3B, C**), indicating that the CHAMP1 complex is required for proper deposition of H3K9 methylation at stalled forks.

**Fig. 3.**
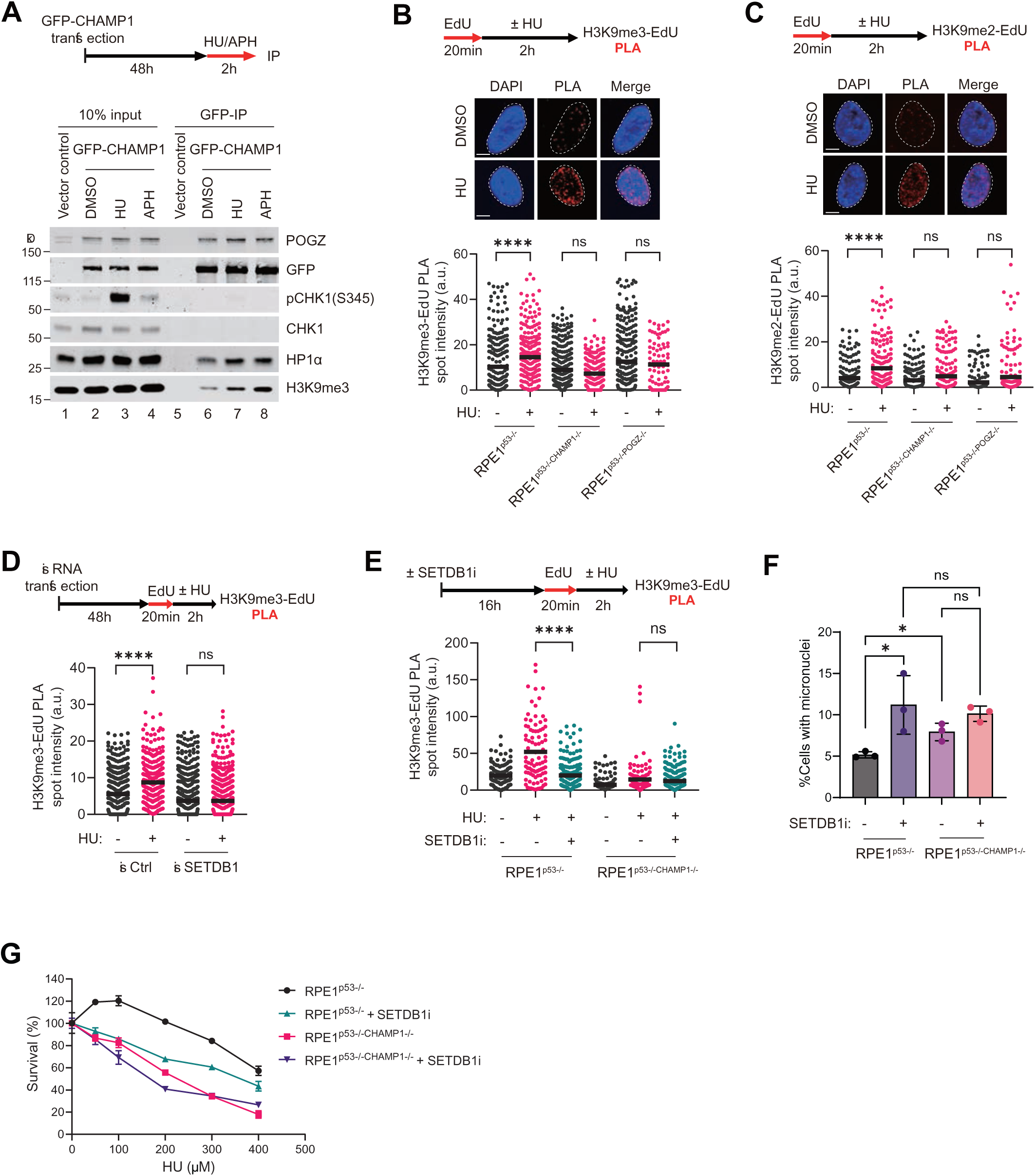
The CHAMP1 complex promotes H3K9me3 accumulation at stalled replication forks. **A.** Western blot analysis of GFP immunoprecipitates from cells expressing either GFP vector control or GFP-CHAMP1, treated with 4 mM hydroxyurea (HU) or 400 nM aphidicolin (APH) for 2 hours. Co-immunoprecipitation of endogenous POGZ, HP1α, and H3K9me3 was assessed. Phosphorylated CHK1 and KAP1 were used as markers of replication stress. Immunoblots are representative of two independent experiments. **B.** Top: Representative images of PLA depicting H3K9me3 presence at replication sites (H3K9me3-EdU PLA, red). Nuclei were stained with DAPI (blue), Scale bar, 5 µm. Cells were treated with EdU for 20 min and were either treated with 4mM HU or DMSO for 2 hours. Bottom: Distribution of the total intensity of all H3K9me3-EdU PLA spots per nucleus. More than 50 cells were counted for each of three independent experiments. P values were calculated using Mann-Whitney test. **C.** Top: Representative images of PLA depicting H3K9me2 presence at replication sites (H3K9me2-EdU PLA, red). Nuclei were stained with DAPI (blue), Scale bar, 5 µm. Cells were treated with EdU for 20 min and were either treated with 4mM HU or DMSO for 2 hours. Bottom: Distribution of the total intensity of all H3K9me2-EdU PLA spots per nucleus. More than 50 cells were counted for each of three independent experiments. P values were calculated using Mann-Whitney test. **D.** RPE1 cells were transfected with siRNA targeting SETDB1 or a non-targeting control (siCtrl) for 48 hours, followed by a 20-minute EdU pulse. Cells were then treated with either 4 mM HU or DMSO for 2 hours. The total intensity of H3K9me3–EdU PLA signals per nucleus was quantified. More than 50 cells were counted for each of three independent experiments. P values were calculated using Mann-Whitney test. **E.** RPE1^p53-/-^ or RPE1^p53-/-CHAMP1-/-^ cells were treated with the SETDB1 inhibitor (SETDB1-TTD-IN-1) for 24 hours, followed by a 20-minute EdU pulse. Cells were then treated with either 4 mM HU or DMSO for 2 hours. The total intensity of H3K9me3–EdU PLA signals per nucleus was quantified. More than 50 cells were counted for each of three independent experiments. P values were calculated using Mann-Whitney test. **F.** Percentage of interphase cells from RPE^p53-/-^ and RPE^p53-/-CHAMP1-/-^ cells containing micronuclei are presented as mean values ±SD, n=3 independent experiments. Cells were treated with SETDB1 inhibitor (SETDB1-TTD-IN-1) or DMSO for 24 hours before imaging. More than 100 cells were counted for each experiment. P values were calculated using a two-tailed Student’s t test. **G.** 3-day cytotoxicity analysis of RPE^p53-/-^ and RPE^p53-/-CHAMP1-/-^ cells treated with various doses of HU, with or without the SETDB1 inhibitor (SETDB1-TTD-IN-1). Cell viability was detected by CellTiter Glo (Promega). Error bars indicate SD, n=3 independent experiments. Cell viability was detected by CellTiter Glo (Promega). Error bars indicate SD, n=3 independent experiments.

To further investigate the mechanism, we tested whether SETDB1 is responsible for *de novo* H3K9me3 deposition at stalled replication forks. Interestingly, depletion of the SETDB1 using siRNA or chemical inhibition of SETDB1 abolished the HU-induced H3K9me3–EdU PLA signals (**Figure 3D, E, Supplemental Figure 3A, B**). Notably, SETDB1 inhibition did not further reduce H3K9me3 levels in CHAMP1 knockout cells, suggesting that SETDB1 may function downstream of CHAMP1. Inhibition of SETDB1 also increased micronuclei formation and sensitized cells to HU; however, these phenotypes that were not further enhanced in CHAMP1-deficient cells (**Figure 3F-G**), suggesting that the CHAMP1 and SETDB1 function epistatically in the response to stalled replication forks. Taken together, our results suggest that the CHAMP1– POGZ complex functions as a key regulator of H3K9me3 deposition at stalled replication forks by coordinating the recruitment and activity of SETDB1 methyltransferase and maintaining replication fork integrity under replication stress.

### CHAMP1 complex promotes ORC recruitment at telomeres and resolves telomeric replication stress in ALT cells

While the CHAMP1 complex appears to play a general role in relieving replication stress, it may have a more specific role in some cellular settings. For instance, previous studies have shown that tumor cells that use the ALT mechanism for telomere maintenance have very high levels of replication stress and homology-directed DNA repair at their telomeres ^14,50,51^ and a high local accumulation of H3K9me3 deposition ^52–55^ ^51^. Indeed, ALT-positive tumor cells rely on several mechanisms to limit their replication stress, such as the upregulation of FANCM and SMARCAL1 activity. ^10^ ^56^. We reasoned that the CHAMP1 complex may also play a role in reducing replication stress in ALT-positive tumors.

To determine whether the CHAMP1 complex is required for ALT telomere replication, we employed PICh (<u>P</u>roteomics of Isolated <u>Ch</u>romatin segments) to profile proteins associated with ALT telomeres in CHAMP1 knockout ALT-positive U2OS cells ^14^ (**Figure 4A**). Strikingly, loss of CHAMP1 markedly reduced the recruitment of origin replication complex (ORC) proteins (**Figure 4B**), which are known to associate with heterochromatin and are critical for telomere replication ^39,41,43,57^. Additionally, CHAMP1 depletion led to elevated levels of FANCM, other Fanconi anemia (FA) proteins, and other replication stress proteins, including ATR, ATRIP, and TOPBP1 (**Figure 4B**) at telomeres. Immunofluorescence further confirmed the reduction of the ORC subunit, ORC2, at telomeres in CHAMP1 knockout and POGZ knockout U2OS cells (**Figure 4C, D; Supplemental Figure 4A, B**). Consistently, depletion of CHAMP1 or POGZ resulted in increased pRPA2 foci enrichment and damage signal at telomeres (**Figure 4E-H**). To further study the function of the CHAMP1 complex for telomere replication, we next performed Telo-FISH (telomere fluorescence in situ hybridization) assays to quantify telomere signals. Remarkably, knockout of CHAMP1 or POGZ significantly increased telomere loss and telomere fragility in U2OS cells (**Figure 4I-K**), known readouts of telomere replication defects ^13^. Collectively, these findings underscore the significance of the CHAMP1 complex in telomere replication and integrity maintenance of ALT cells.

**Fig. 4.**
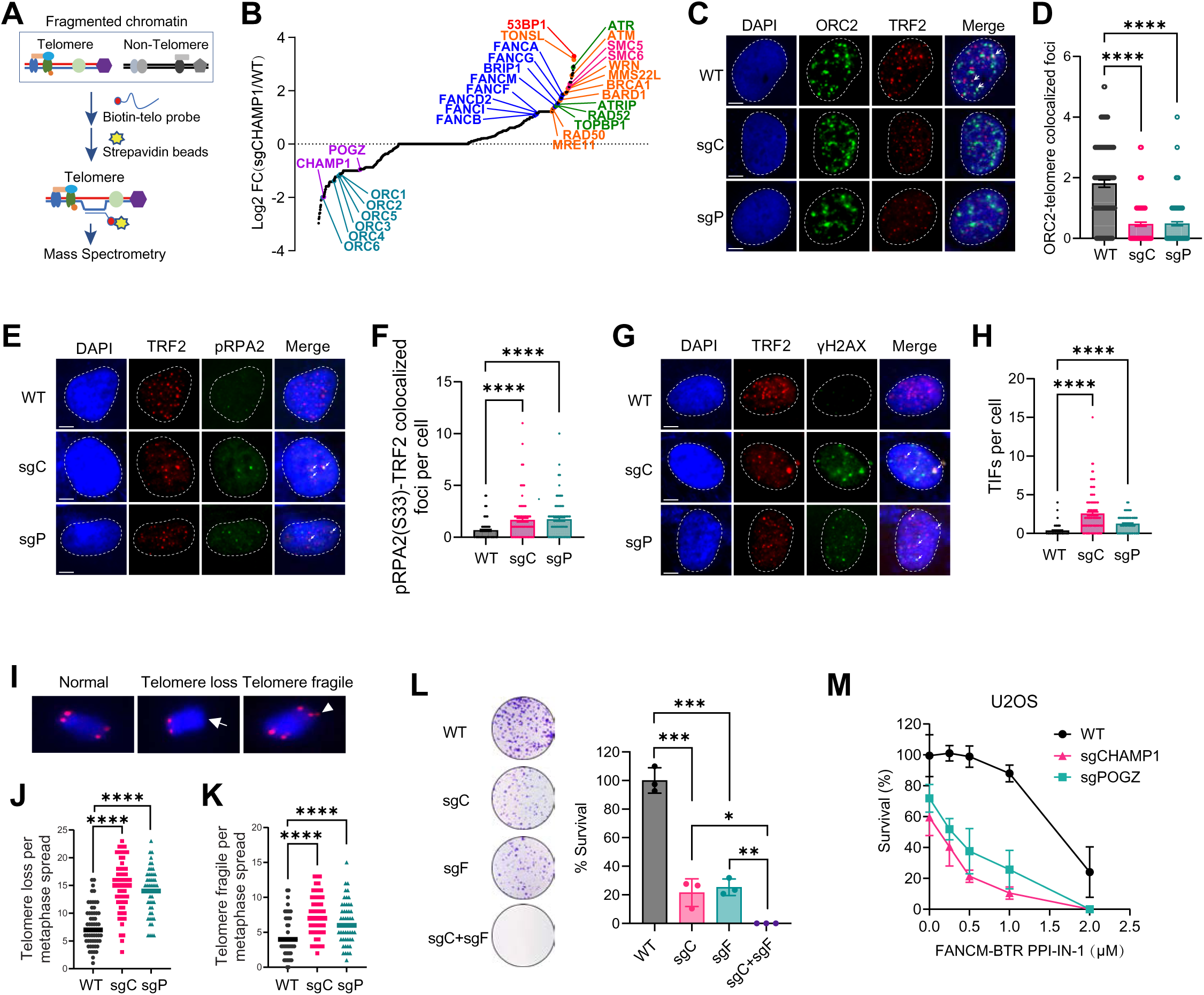
The CHAMP1 complex reduces telomere replication stress and promotes ORC recruitment in ALT cells. **A.** Schematic of the PICh (Proteomics of Isolated Chromatin segments) strategy used to profile the telomere-associated proteome in WT and sgCHAMP1 U2OS cells. Chromatin was fractionated and pre-cleared, then hybridized with a biotinylated telomeric probe and captured using magnetic beads. Telomere-associated proteins were subsequently identified by mass spectrometry. **B.** Scatterplot showing the Log₂(sgCHAMP1/ WT) values of total peptide counts from telomere-enriched proteomes in WT and sgCHAMP1 U2OS cells. Replication stress response proteins and ORC complex components are highlighted. **C.** Representative confocal images of ORC2 colocalization with TRF2 (telomere marker protein) in WT, sgCHAMP1 (sgC) and sgPOGZ (sgP) U2OS cells. DAPI was used to stain the nuclei. Scale bar, 5 μm. **D.** Quantification of ORC2-TRF2 colocalizations in (i). Data are presented as mean values ±SEM. More than 50 cells were counted for each of three independent experiments. P values were calculated using Mann-Whitney test. **E.** Representative confocal images of pRPA2(S33) colocalization with TRF2 in WT, sgCHAMP1 (sgC) and sgPOGZ (sgP) U2OS cells. DAPI was used to stain the nuclei. Scale bar, 5 μm. **F.** Quantification of pRPA2-TRF2 colocalizations in (e). Data are presented as mean values ±SEM. More than 50 cells were counted for each of three independent experiments. P values were calculated using Mann-Whitney test. **G.** Representative confocal images of γH2AX colocalization with TRF2 in WT, sgCHAMP1 (sgC) and sgPOGZ (sgP) U2OS cells. DAPI was used to stain the nuclei. Scale bar, 5 μm. **H.** Quantification of γH2AX-TRF2 colocalizations in (g). Data are presented as mean values ±SEM. More than 50 cells were counted for each of three independent experiments. P values were calculated using Mann-Whitney test. **I.** Representative images of chromosomes showing normal telomeres, telomere loss (arrow), and fragile telomeres (arrowhead). **J–K.** Quantification of telomere loss (**J**) and telomere fragility (**K**) per metaphase spread in WT, CHAMP1-knockout (sgCHAMP1), and POGZ-knockout (sgPOGZ) U2OS cells. **L.** Colony survival assay of WT and sgCHAMP1 (sgC) U2OS cells treated with sgFANCM (sgF). **M.** A10-day clonogenic survival of WT, sgCHAMP1 and sgPOGZ U2OS cells treated with various dose of FANCMi (FANCM-BTR PPI-IN-1).

The ATR and FANCM proteins are highly enriched in ALT-positive telomeres and may function to alleviate replication stress and to maintain tumor cell viability ^8,10,11,58,59^. Given the critical role of FANCM in maintaining genome stability in ALT-positive cells, FANCM inhibitors have emerged as promising therapeutic agents for the effective suppression of ALT-driven tumors. Remarkably, simultaneous knockout of CHAMP1 and FANCM induces synthetic lethality in U2OS cells (**Figure 4L**). Moreover, ALT cells deficient in CHAMP1 complex subunits exhibit significantly increased sensitivity to a FANCM inhibitor (**Figure 4M; Supplemental Figure 4C**), highlighting a potential novel synthetic lethal strategy for targeting ALT-associated cancers. Taken together, knockout of the CHAMP1 complex increases telomere replication stress and sensitizes the ALT cells to drugs which further increase replication stress.

### CHAMP1 complex orchestrates H3K9me3-dependent ORC recruitment to facilitate replication

To further determine whether the CHAMP1 complex plays a more general role in the loading of ORC at sites of replication stress beyond its role in ALT-positive cells, we examined ORC2 recruitment at stalled replication forks. In wild-type cells, ORC2 was enriched at HU-stalled forks, whereas this enrichment was markedly reduced in CHAMP1-deficient cells (**Figure 5A, B**). Importantly, treatment with a SETDB1 inhibitor also abolished ORC2 accumulation at stalled forks (**Figure 5A, B**), suggesting that H3K9me3 is required for ORC2 recruitment under replication stress. This finding is consistent with previous reports showing preferential ORC binding to heterochromatic regions ^43,57,60^. To further explore the functional relevance of ORC2 at stalled replication forks and its regulation by CHAMP1, we performed siRNA-mediated knockdown of ORC2 in CHAMP1 knockout cells (**Supplemental Figure 5**). ORC2 depletion sensitized cells to HU and aphidicolin treatment (**Figure 5C, D**), phenocopying the effect of CHAMP1 loss. Notably, co-depletion of CHAMP1 and ORC2 did not further increase sensitivity (**Figure 5C, D**), suggesting that they function in the same pathway. In summary, CHAMP1 facilitates ORC2 recruitment to stalled replication forks by promoting H3K9me3 accumulation, thereby preserving fork stability under replication stress.

**Fig. 5.**
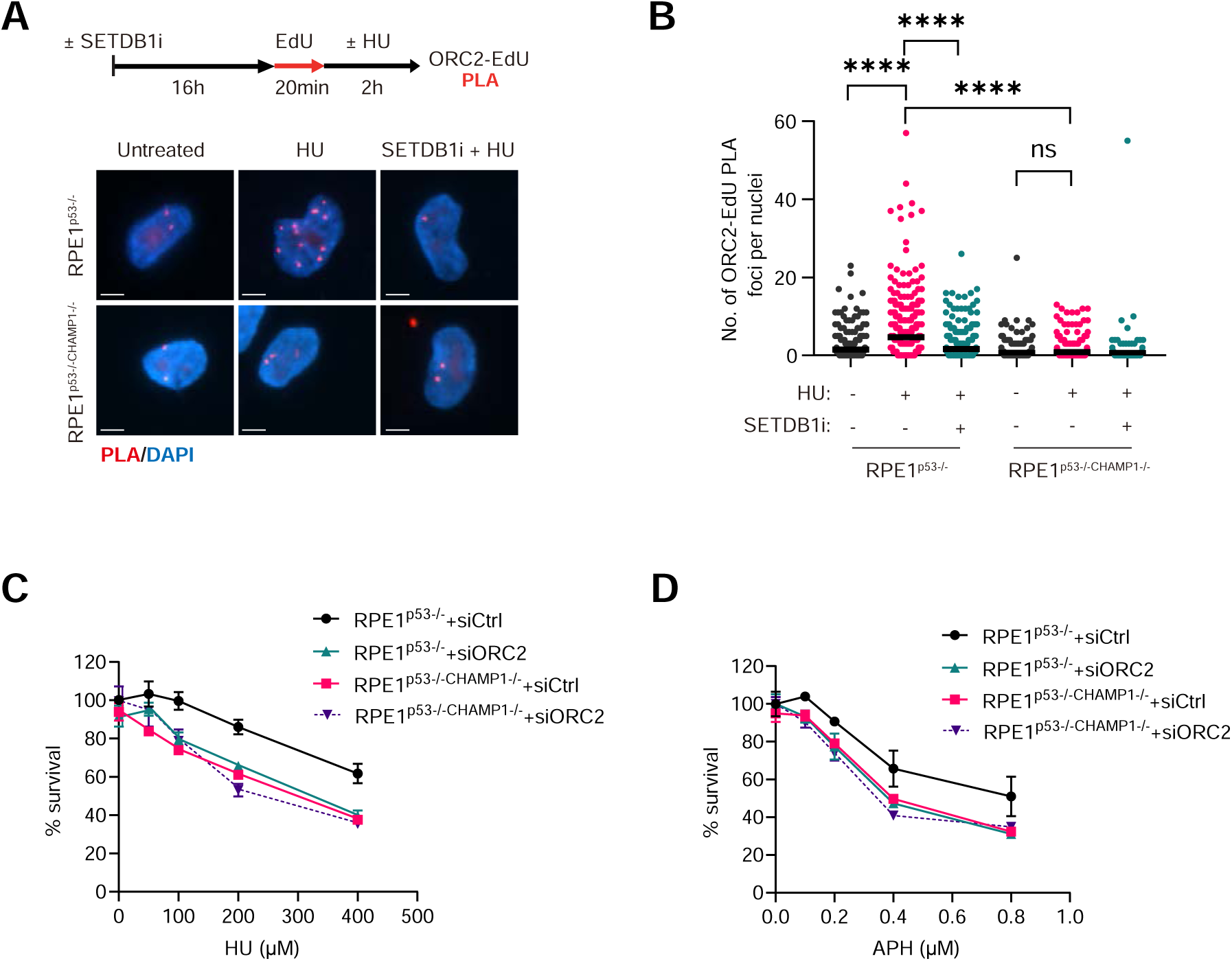
The CHAMP1 complex orchestrates H3K9me3-dependent ORC recruitment to facilitate replication. **A.** Representative images of PLA depecting ORC2 presence at replication sites (ORC2-EdU PLA, red). Nuclei were stained with DAPI (blue), Scale bar, 5 µm. **B.** RPE1^p53-/-^ or RPE1^p53-/-CHAMP1-/-^ cells were treated with the SETDB1 inhibitor for 24 hours, followed by a 20-minute EdU pulse. Cells were then treated with either 4 mM HU or DMSO for 2 hours. The total intensity of H3K9me3–EdU PLA signals per nucleus was quantified. More than 50 cells were counted for each of three independent experiments. P values were calculated using Mann-Whitney test. **C.** Western blot analysis of GFP immunoprecipitates from cells expressing either GFP vector control or GFP-CHAMP1, treated with 4 mM hydroxyurea (HU) or 400 nM aphidicolin (APH) for 2 hours. Co-immunoprecipitation of endogenous ORC2 was assessed. Immunoblots are representative of two independent experiments. **D.** Western blot analysis of GFP immunoprecipitates from cells expressing either GFP vector control or GFP-CHAMP1. Cells were treated with or without SETDB1 inhibitor for 24 hours, followed by treatment with 4 mM hydroxyurea (HU) for 2 hours. Co-immunoprecipitation of endogenous ORC2 was assessed. Immunoblots are representative of two independent experiments. **E.** Immunoblotting against CHAMP1 and ORC2 in RPE1^p53-/-^ and RPE1^p53-/-CHAMP1-/-^ cells after 48h siRNA treatment against ORC2. GAPDH was used as a loading control. **F.** DNA fiber assay was performed using the same cells as in (E) to assess replication fork degradation. Cells were sequentially labeled with CIdU and IdU for 45 minutes each, followed by treatment with 4 mM HU or DMSO for 4 hours. Fork degradation was quantified as the ratio of IdU to CIdU tract lengths. Each dot represents an individual fiber; at least 200 fibers were analyzed per condition. Statistical significance was assessed using the Mann–Whitney U test. **G.** Replication fork restart was analyzed in the same cells as in (E) using DNA fiber assay. Cells were labeled with CIdU, treated with 4 mM HU for 1 hour, then released into IdU for 45 minutes. Fork restart was quantified by measuring IdU tract lengths. Each dot represents an individual fiber; at least 200 fibers were analyzed per condition. Horizontal bars represent medians. Statistical significance was determined using the Mann–Whitney U test. **H-I.** 3-day cytotoxicity analysis of RPE1^p53-/-^ and RPE1^p53-/-CHAMP1-/-^ cells treated with various doses of HU (**H**) or aphidicolin (APH) (**I**) after 48h siCtrl or siORC2 treatment. Cell viability was detected by CellTiterGlo (Promega). Error bars indicate SD, n=3 independent experiments.

### Cancer cells under replication stress dependent on the CHAMP1 complex

High-grade ovarian cancer and many cancers undergo replication stress due to amplification of oncogene cyclin E1 (CCNE1) ^47^. To study how tumors survive under elevated replication stress, we performed gene expression and dependency analyses on breast and ovarian cancer cells with CCNE1 amplification. Consistent with previous findings, DepMap data analysis revealed that FA complex mRNA levels strongly correlate with CCNE1 mRNA expression (**Figure 6A**). Interestingly, DepMap data analysis also revealed that CHAMP1 complex and ORC complex mRNA levels strongly correlate with CCNE1 mRNA expression (**Figure 6A**). Consistently, protein analysis confirmed high CHAMP1 levels in CCNE1-amplified ovarian cancer cells (**Figure 6B, C**). Importantly, CHAMP1 depletion selectively impaired viability in CCNE1-amplified cancer cell lines (**Figure 6D; Supplemental Figure 6**). Moreover, TCGA analysis showed that high CHAMP1 expression predicted worse survival in patients with CCNE1-amplified tumors, but not in those without amplification (**Figure 6E, F**). These results highlight a potential oncogene-induced dependency on the CHAMP1 complex, revealing therapeutic opportunities in cancers driven by replication stress.

**Fig. 6.**
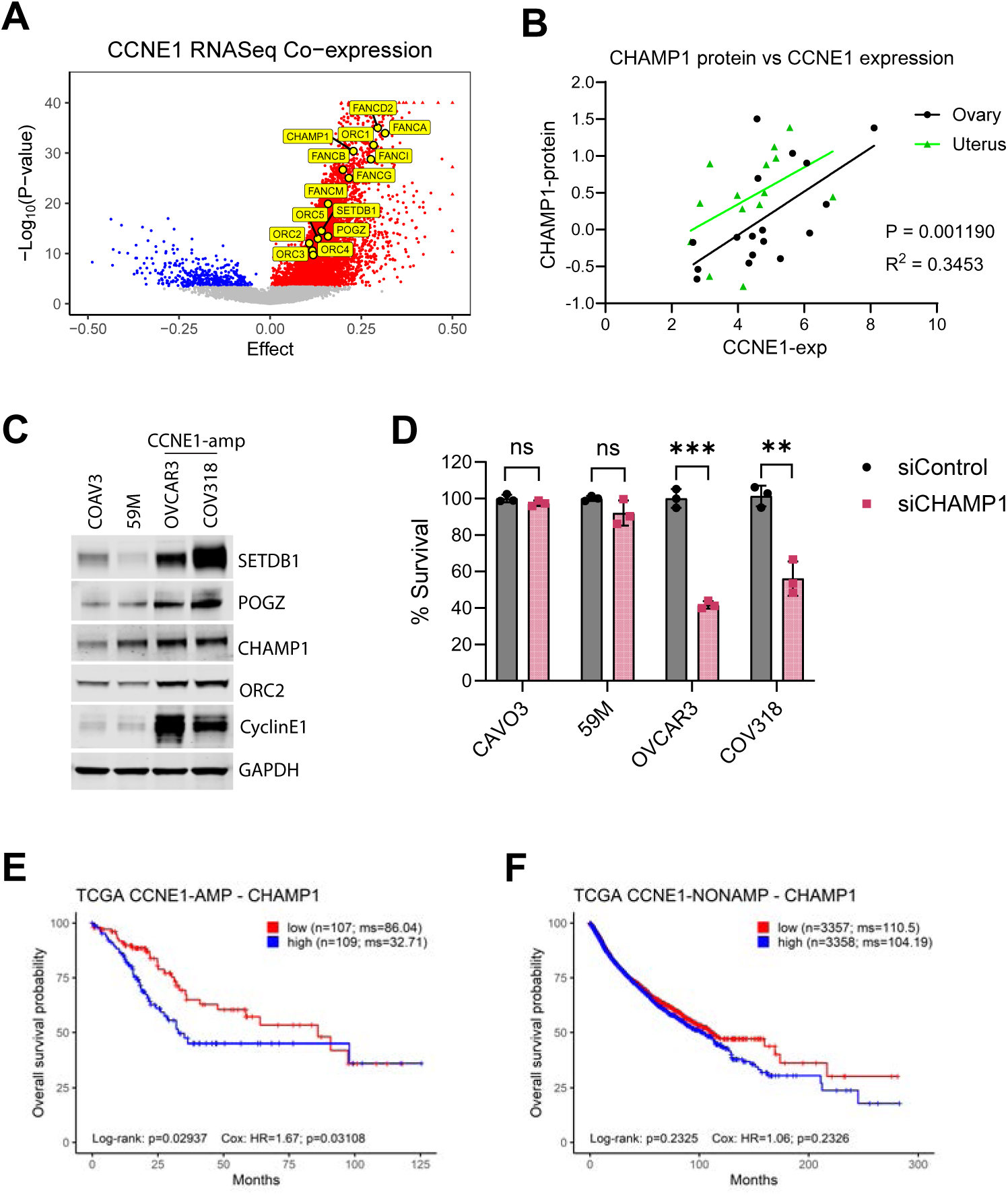
Cancer cells with replication stress are highly dependent on CHAMP1. **A.** Volcano plot showing the genes co-expressed with CCNE1. Replication stress response proteins, CHAMP1 complex, and ORC complex were highlighted. **B.** CHAMP1 protein levels positively correlate with cyclin E mRNA expression in ovarian and uterine cancer cell lines. **C.** Immunoblot analysis of ovarian cancer cell lines with or without *CCNE1* amplification (amp), using the indicated antibodies. GAPDH acts as a loading control. **D.** Survival assay of ovarian cancer cell lines treated with siControl or siCHAMP1 for three days. **E-F.** Kaplan–Meier survival curves showing overall survival of patients from TCGA with high CHAMP1 expression, stratified by *CCNE1* amplification status: *CCNE1*-amplified (**E**) and non-amplified (**F**). This analysis combines tumors of all cancer types from TCGA studies. N represents the number of patients.

## DISCUSSION

Our recent study indicates that the CHAMP1 complex plays a role in the assembly of heterochromatin at multiple genomic sites ^13^. The CHAMP1 complex binds and recruits the methyltransferase SETDB1, leading to H3K9me3 deposition, which in turn facilitates telomere clustering and promotes homologous recombination (HR) at these sites ^13^. We now demonstrate that the CHAMP1 complex plays a more general role in the reduction of cellular replication stress. Accordingly, loss of CHAMP1 or its direct interactor POGZ results in basal and hydroxyurea-inducible single-strand DNA accumulation, a hallmark of replication stress. Deleting CHAMP1 led to an increase in the activation of the DNA damage checkpoint ATR/CHK1 pathway^47^. CHAMP1-deficient cells are hypersensitive to inhibitors of the ATR/CHK1 pathway and the Fanconi Anemia (FA) pathway.

Mechanistically, we demonstrate that the CHAMP1 complex safeguards stalled replication forks through several coordinated mechanisms (**Figure 7**). First, it promotes the deposition of H3K9me3 via recruitment and activation of SETDB1, establishing a repressive chromatin environment that stabilizes the fork. Second, it limits excessive DNA end resection by blocking recruitment of the MRE11 nuclease, thereby protecting nascent DNA from degradation. Third, it facilitates the recruitment of ORC, which may contribute to replication fork stability or restart, although the precise mechanism remains to be fully elucidated. Together, these activities preserve fork integrity and maintain genome stability under conditions of replication stress.

**Fig. 7.**
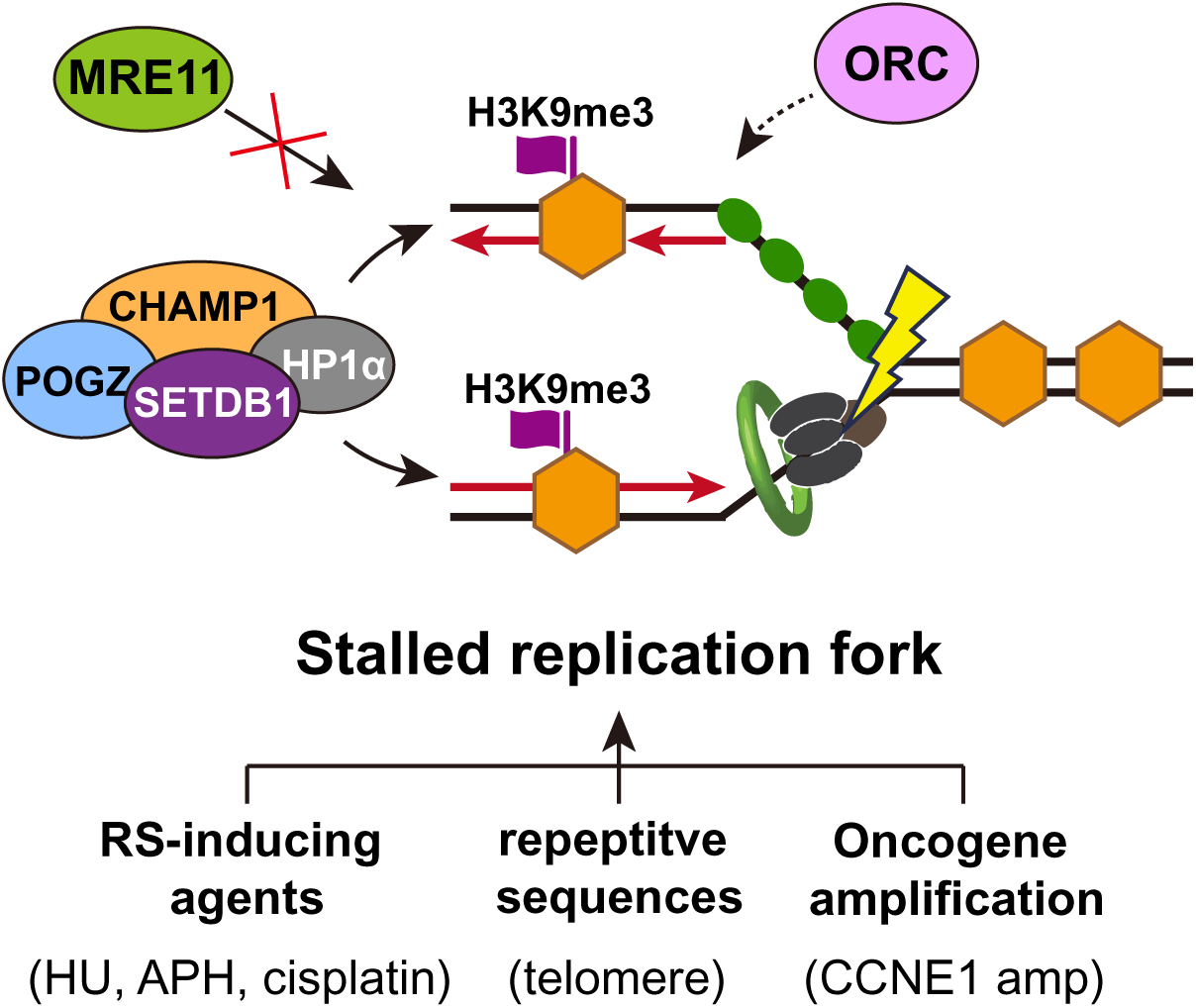
Schematic model illustrating how the CHAMP1 complex regulates stalled replication fork stability. The CHAMP1 complex—comprising CHAMP1, POGZ, HP1α, and SETDB1—plays a critical role in maintaining the stability of stalled replication forks under replication stress, which can arise from repetitive DNA sequences, oncogene deregulation, or exogenous agents. This complex functions through multiple coordinated mechanisms: (1) it promotes the deposition of H3K9me3 via recruitment and activation of the SETDB1 methyltransferase, establishing a heterochromatic environment at stalled forks; (2) it prevents excessive resection by inhibiting the recruitment of the MRE11 nuclease, thereby protecting nascent DNA from degradation; and (3) it facilitates the recruitment of the ORC complex, potentially promoting replication fork restart. Together, these actions ensure replication fork integrity and safeguard genome stability during conditions of replication stress.

Previous study has shown that H3K9me3 enrichment at stalled replication forks is also mediated by G9a and is critical for maintaining fork integrity ^49^. It suggests that multiple H3K9 methyltransferases may coordinate to modulate heterochromatin formation at stressed forks. Future studies are needed to elucidate how SETDB1 and G9a dynamically regulate H3K9 methylation at stalled replication forks, and how their coordinated activity—potentially orchestrated by CHAMP1—contributes to fork stabilization and genome integrity.

Telomeres maintained by the ALT pathway exhibit elevated replication stress ^51^. Loss of CHAMP1 further elevates replication stress at these sites, as evidenced by increased recruitment of proteins from the ATR/CHK1 and FA pathways, including FANCM ^61^. These cells also show hypersensitivity to inhibitors of ATR or FANCM, highlighting a functional dependency on these replication stress response mechanisms when CHAMP1 function is compromised.

A particularly novel finding from our work is the requirement of CHAMP1 for the recruitment of the Origin Recognition Complex (ORC) to ALT telomeres and stalled replication forks. ORC is best known for its role in replication origin licensing, and has also been shown to localize to telomeres via interactions with TRF2 ^41,42^ and HP1 ^43^. Our data suggest that CHAMP1 is a previously unrecognized upstream regulator of ORC loading at telomeres and stalled replication forks, although whether this occurs via direct physical interaction or through chromatin remodeling remains to be determined. Deletion of ORC2 led to increased sensitivity to hydroxyurea, suggesting that ORC plays a role in the replication stress response. Future studies are needed to define the mechanistic contribution of ORC to stalled fork processing and restart, particularly in the context of heterochromatinized regions. It will also be important to determine how CHAMP1-mediated ORC recruitment integrates with other fork protection pathways to ensure genome stability.

Beyond cellular models, our findings have important clinical implications. Humans with *de novo* heterozygous mutations in *CHAMP1* exhibit a neurodevelopmental disorder, the CHAMP1 syndrome, characterized by intellectual disability, behavioral symptoms, and distinct dysmorphic features ^22–26^. A recent study indicates that rare patients with CHAMP1 syndrome develop leukemia, suggesting a broader role of the CHAMP1 complex in limiting replication stress and maintaining the genomic stability of hematopoietic stem and progenitor cells ^22^. *De novo* heterozygous mutations in the *POGZ* gene also result in a rare but highly-related neurodevelopment syndrome called the White Sutton Syndrome (WSS) ^27–35^. Using EBV-transformed peripheral blood lymphoblasts from patients with CHAMP1 syndrome, we have demonstrated a cellular defect in heterochromatinization and homologous recombination, based on the reduction in RAD51 foci assembly. These patient-derived lymphoblastoid cell lines (LCLs) also display marked HU hypersensitivity, reinforcing the conclusion that CHAMP1 deficiency leads to elevated baseline RS. Whether LCLs from WSS patients have similar replication stress and HU sensitivity remains to be determined.

Replication stress in tumor cells can result from several mechanisms, including loss of the p53 protein, mitogenic enhancement from oncoprotein expression, or disruption of DNA repair pathways. Cancer cells with replication stress upregulate the ATR/CHK1 pathway as a mechanism to enhance cell cycle checkpoints and to slow down cell growth. We reasoned that upregulation of the CHAMP1 complex might provide an independent mechanism for allowing cancer cells to tolerate their replication stress. Consistent with this hypothesis, we have shown that cancer cells that are driven by CCNE1 amplification not only exhibit upregulated expression of CHAMP1 and POGZ but are also functionally dependent on these factors for survival under replication stress conditions.

In summary, the CHAMP1 complex plays a broad role in the cellular response to stalled forks by promoting H3K9me3 enrichment via SETDB1 and preventing nascent DNA degradation through inhibition of MRE11 recruitment. CHAMP1 also facilitates the recruitment of ORC to sites of replication stress, including ALT telomeres and HU-stalled forks. Disruption of the CHAMP1 complex, or inhibition of its associated methyltransferase SETDB1, may offer a therapeutic strategy to exacerbate replication stress and genomic instability in cancer cells, thereby selectively eliminating tumor cells. Moreover, as CHAMP1 and POGZ mutations are also associated with neurodevelopmental disorders, further studies into their functions may provide broader insights into both tumorigenesis and human genetic disease.

## MATERIALS AND METHODS

### Cell Culture Conditions

Human U2OS and RPE1-hTERT cells were cultured in DMEM/F12 + Glutamax (Invitrogen) supplemented with 10% FBS (Sigma) and 1% penicillin-streptomycin (Invitrogen). 293T cells were grown in DMEM (Thermo Fisher) with 10% FBS and 1% penicillin-streptomycin. Each of these cell lines were obtained from the American Type Culture Collection (ATCC) and validated by short tandem repeat (STR) testing. CHAMP1 patient lymphocyte cell lines from Coriell Institute were grown in RPMI1640 with 15% FBS (sigma) and 1% penicillin-streptomycin (Thermo Fisher). Cell lines were maintained in an incubator at 37 °C and 5% CO2 according to standard protocols. Cell lines were validated to be negative for mycoplasma contamination using the MycoAlert Plus Mycoplasma Detection Kit (Lonza).

### RNA interference and CRISPR-mediated gene deletions

siRNA knockdown and CRISPR knockout were carried out as previously described ^13^. siRNA knockdown experiments were performed out using RNAiMax (Invitrogen, #13778150) according to the manufacturer’s protocols. See Key Resource Table for siRNAs used. For single-gene knockout, sgRNAs targeting candidate genes were either cloned into the pSpCas9 BB-2A-GFP (PX458) vector (GenScript) or delivered into cells together with Cas9 protein via electroporation (Lonza), according to the manufacturer’s protocol. See Key Resource Table for the sgRNA sequences used. For PX458-mediated transfections, GFP-positive cells were sorted 48 hours post-transfection using a BD FACSAria II cell sorter. Knockdown and knockout efficiency were validated by western blotting.

### Proximity Ligation Assay

Proximity ligation assay was carried out as previously described ^62^. Cells were grown on 10mm coverslips in 24-well dishes until 60% confluence. SETDB1 inhibitors were added for 16 h. Cells were grown with EdU (5-ethynyl-2’-deoxyuridine) (Invitrogen, A10044) (20 µM) for 10 minutes to label nascent DNA strands. Cells were then grown in hydroxyurea (4 mM) for 2 h to induce stalled replication forks. Coverslips were washed with 1X PBS followed by 0.5 Triton X-100 for 5 minutes. Cells were fixed in 4% paraformaldehyde for 10 minutes at room temperature. Cells were then washed twice with 1X PBS and then in blocking buffer 3% BSA in 0.1 Triton X-100 for 1 h at room temperature. Following blocking, cells were incubated in click reaction mix (1X PBS, Copper (II) sulfate pentahydrate (5 mM) (Sigma Aldrich, #203165-10G), Biotin Azide (PEG4 carboxamide-6-Azidohexanyl Biotin) (10 μM) (Life Technologies, #B10184), and sodium ascorbate (10 mM) (Santa Cruz Biotechnology, #SC-215877)) for 30 minutes protected from light. Coverslips were incubated with desired primary antibodies overnight at 4C°: mouse α-biotin (Thermo Fisher Scientific, #NC9724635) (1:1000 dilution), rabbit α-H3K9me3 (Abcam, #ab8898-100ug) (1:1000 dilution), rabbit α-H3K9me2 (Abcam, #ab1220), and rabbit α-ORC2 (Life Technologies, # PA5-70227) (1:500 dilution).

Following primary antibody incubation, coverslips were incubated with PLA probes Duolink In Situ PLA Probe Anti-Rabbit PLUS (DUO92002) and Duolink In Situ PLA Probe Anti-Mouse MINUS (DUO92004) as in the provided instructions. Duolink In situ detection kit (DUO92008) was used for the ligation and amplification of the PLA signal. Coverslips were mounted with Prolong diamond antifade reagent with DAPI (ThermoFisher Scientific, P36971). Images were captured on a Zeiss microscope and analyzed with Cell Profiler.

### Immunofluorescence assay

Cells were seeded onto glass coverslips placed in 24-well plates. For immunofluorescence, the cells were pre-extracted with 0.5% Triton X-100 for 5 minutes, followed by fixation with 4% paraformaldehyde for 10 minutes at 4°C. After three PBS washes, a blocking step was carried out using 3% BSA in PBS for 1 hour at room temperature. Coverslips were then incubated with primary antibodies overnight at 4°C, followed by incubation with secondary antibodies for 1 hour at room temperature. Finally, coverslips were mounted with DAPI (Vector Laboratories) and imaged using a Zeiss AX10 fluorescence microscope with Zen software.

### Immunoprecipitation

After GFP-CHAMP1 plasmid transfection for 48 hours, 293T cells were treated with HU, APH or DMSO treatment for 2hours. 293T cells were subsequently collected and subjected to lysis using NETN lysis buffer containing a proteinase and phosphatase inhibitor cocktail (Thermo, 1:100) for 30 minutes on ice. Following this, the lysed samples were incubated overnight at 4°C with GFP-Trap_A (Chromotek). The beads were then thoroughly washed four times with NETN buffer, and the immunoprecipitated materials were eluted by boiling. Western blot analysis was conducted to detect the immunoprecipitates.

### Western Blot Analysis

Cells were lysed with 1X RIPA buffer (Cell Signaling Technology, #9806) with protease and phosphatase inhibitors on ice for 10 minutes. Cell lysate was centrifuged at 15,000 rpm for 10 min at 4C°. Supernatant was collected and mixed with equal volume of 2x Laemmli sample buffer (BioRad #1610737) and boiled at 100C° for 10 minutes. Lysates were resolved on precast gels and transferred onto nitrocellulose membranes. Membranes were blocked with 5% milk in TBST and probed with primary and secondary antibodies respectively and then detected by immunofluorescence with an ODYSSEY CLx machine using ImageStudio Ver. 5.2.

### Telomere-FISH

Quantitative telomere FISH (Fluorescence In Situ Hybridization) was performed as previously described ^63^. Briefly, cells were synchronized using nocodazole (0.5mg/ml) treatment for 4–6 hours, then collected and incubated in 0.075 M KCl at 37°C for 25 minutes. After hypotonic treatment, cells were fixed in methanol/acetic acid (3:1) at -20°C overnight and then spread onto slides. Telomere FISH was conducted using a TelG-Cy3 PNA probe (PNAbio) according to established protocols.

### Clonogenic Assays

Cells were counted and plated in triplicate in 6-cell plates for colony formation assays. After 24 h, various drugs were added at various concentrations and cells were incubated for 10 days. Cells were then fixed with 4% formaldehyde for 10 min. at room temperature and then stained with crystal violet for 30 minutes. Cell growth area quantification and imaging of the plates were performed with GelCount (Oxford Optronix).

### Cell-Titer-Glo Luminescent Cell Viability Assay

Short-term CellTiter-Glo survival assays were conducted as previously described ^64^. To assess drug sensitivity, cells were seeded in 96-well plates at a density of 500 cells per well. After 12 hours, cells were treated with the indicated concentrations of drug. Cell viability was measured three days later using the CellTiter-Glo assay (Promega). Survival at each drug concentration was calculated as a percentage relative to the corresponding untreated control.

### Proteomics of Isolated Chromatin segments (PICh)

PICh assay was carried out as previously described ^13^.

### Fork Stability and Restart Assay

To detect fork stability and restart we used DNA fiber assays ^62^. For fork stability assay, cells were pulse-labeled with 100 µM CldU (5-Chloro-2′-deoxyuridine, Sigma C6891) for 45 min, followed by 100 µM IdU (5-Iodo-2′-deoxyuridine, Sigma I7125) for 45 min. Forks were stalled by treating the cells with HU (4mM) for 4 h before harvesting. For fork restart assay, cells were first labeled with CldU (100 µM, 45 min), then treated with HU (4 mM, 1 h) to stall replication forks. After HU washout, fork restart was monitored by labeling with IdU (100 µM, 45 min). Fiber assay was performed as per FiberPrep DNA extraction protocol (Genomic Vision, cat# EXT001A). Briefly, cells were resuspended in PBS and embedded in agarose plugs, which were subsequently treated with proteinase K overnight. Agarose plugs were then digested with beta-agarase overnight. Samples were then poured into FiberComb wells and combed onto silanized coverslips (Genomic Vision, cat# COV-002) using the Molecular Combing System from Genomic Vision. Following denaturation, coverslips were probed with rat anti-BrdU antibody (clone BU1/75 (ICR1) specific to BrdU, Life Technologies MA182088) and mouse anti-IdU Antibody (specific to IdU, BD Biosciences 347580) overnight at 37°C. Next day, the coverslips were washed with PBS-T (0.05% Tween) and incubated with secondary goat anti rat Cy5 antibody (Abcam cat# ab6565) and goat anti mouse Cy3 antibody (Abcam cat #97035) for 45 minutes at 37°C, followed by mouse anti-ssDNA antibody (cat# DSHB auto anti-ssdna) for 1 hour and 15 minutes at 37°C, and goat anti mouse BV480 (Jackson cat #115-685-166) for 45 minutes at 37°C. The coverslips were mounted, and the fibers were scanned using FiberVision S. The lengths of IdU tracts or IdU/CldU ratio were measured by ImageJ and graphed using Prism9 (GraphPad).

### Cancer Cell Line Acquisition Data and Analysis

Co-expressed genes of CCNE1 were determined by assessing the associations between its mRNA expression and that of all other genes. The log₂(TPM+1) – normalized mRNA expression data of the human cancer cell lines from the Cancer Cell Line Encyclopedia (CCLE) ^65^ project were downloaded via the Broad Institute’s Cancer Dependency Map (DepMap; release 19Q4 Public). The associations between mRNA expression of CCNE and that of every other gene were assessed using the robust linear regressions, as computed by the limma ^66^ package in R, with robust empirical Bayes moderation and lineage correction. Volcano plot of the results was generated using the ggplot2 package in R.

The association between CHAMP1 protein expression and CCNE1 mRNA expression was assessed using the multiple linear regression with lineage correction, as computed by GraphPad Prism (version 10.4.2). Relative protein expression data for CHAMP1 and the log₂(TPM+1) – normalized mRNA expression data of CCNE1 in the human cancer cell lines were downloaded from DepMap (release 24Q4 Public).

### TCGA Data Acquisition and Analysis

The survival analyses of the TCGA cancer patients were performed using the clinical, CCNE1 amplification data and CHAMP1 mRNA expression data of the TCGA Pan-Cancer study, downloaded from the cBioPortal for Cancer Genomics (cbioportal.org). Patients were stratified into low and high CHAMP1 expression groups based on the median mRNA expression. Survival analyses were performed in R, separately for tumors with and without CCNE1 amplification, to determine whether there was a difference in the overall survival between the two CHAMP1 groups. Kaplan-Meier curves were created, and the log-rank test was used to test for a difference in overall survival using the survival (version 3.8-3) package in R. The p values were calculated from the chi-square distribution. The survminer (version 0.5.0) R package was used to estimate median survivals, and to plot the Kaplan-Meier curves. Additionally, Cox proportional hazards regressions were performed to estimate the hazard ratios between the low- and the high-CHAMP1 groups.

### Quantification and Statistical Analysis

All values are expressed as standard deviation (SD) or standard error of the mean (SEM) as indicated in figure legends. The statistical significance of differences was assessed by Student’s t-test for comparison of two groups and one-way analyses of variance (ANOVA) with Mann-Whitney U test for comparison of multiple groups using Graphpad Prism 9 and 10.

## Acknowledgements

We thank all members of the D’Andrea laboratory for their helpful suggestions and comments. This work was supported by grants from the US National Institutes of Health (R01HL052725), the Breast Cancer Research Foundation, the Ludwig Center at Harvard, the Smith Family Foundation, and the Innovations Research Fund of the Dana-Farber Cancer Institute (A.D.D.). This work was also supported by the Claudia Adams Barr Program in Innovative Basic Cancer Research (F.L and A.S).

## Author contributions

F.L. and A.D.D. conceived the study, analyzed the data, and wrote the manuscript. A.E., F.Z., T.Z., R.R., L.S. performed experiments and analyzed the data. H.N. performed the computational analysis and analyzed the data.

