## Supplemental figure for "CHAMP1 Complex Promotes Heterochromatin Assembly and Reduces Replication Stress"

### Supplemental Figure 1

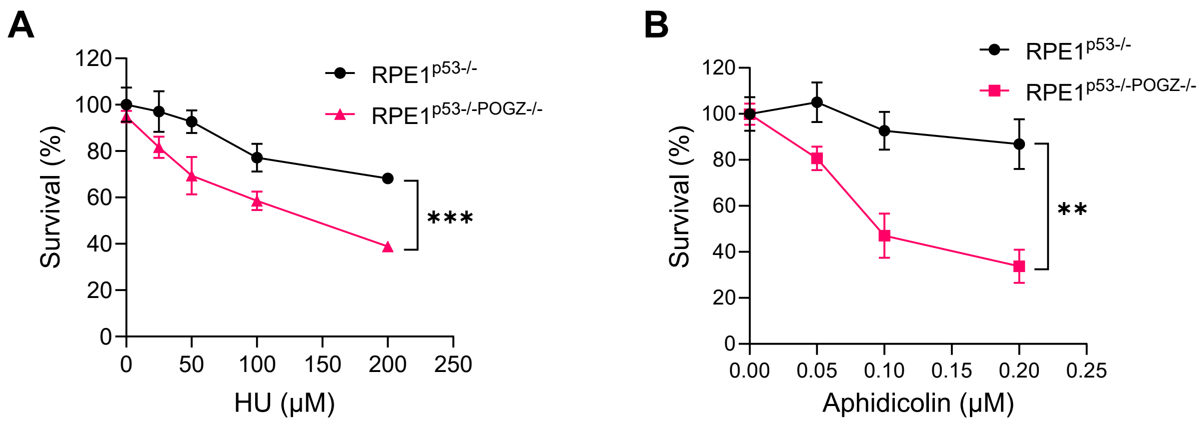


**Supplemental Figure 1, related to Fig. 1. Loss of POGZ enhances cellular sensitivity to replication stress–inducing agents. A-B.** Colony survival plots of RPE1$^{p53-/-}$ and RPE1$^{p53-/-POGZ-/-}$ cells treated with different concentrations of hydroxyurea (HU) (**A**) or aphidicolin (**B**) for 10 days.

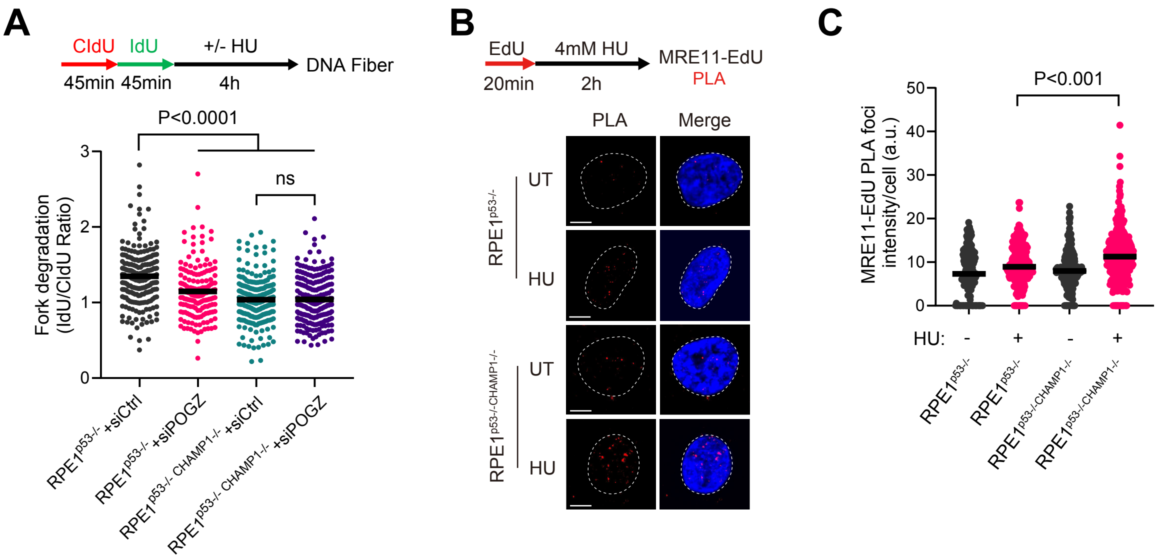


**Supplemental Figure 2, related to Fig. 2. The CHAMP1 complex promotes replication fork stability. A.** Top: Cells were sequentially labeled with CIdU and IdU for 45 minutes each, followed by treatment with either 4 mM HU or DMSO for 4 hours before performing the DNA fiber assay. Bottom: Quantification of replication fork degradation, shown as the ratio of IdU to CIdU tract lengths. Each dot represents an individual fiber, with at least 200 fibers analyzed per condition. Statistical significance was determined using the Mann–Whitney U test. **B.** Representative images of PLA depecting MRE11 presence at replication sites (MRE11-EdU PLA, red). Nuclei were stained with DAPI (blue). Cells were treated with EdU for 20 min and were either treated with 4mM HU or DMSO for 2 hours. **C.** Distribution of the total intensity of all MRE11-EdU PLA spots per nucleus. More than 50 cells were counted for each of three independent experiments. P values were calculated using Mann-Whitney test.

#### Supplemental Figure 3

A

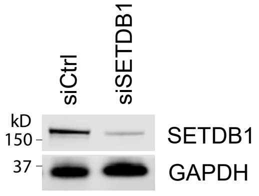


B

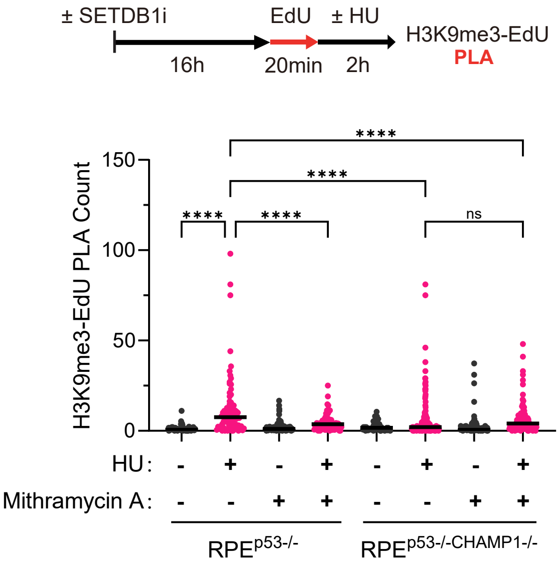


**Supplemental Figure 3, related to Fig. 3. CHAMP1 recruits SETDB1 to deposit H3K9me3 at stalled replication forks. A.** Immunoblotting against SETDB1 in RPE1$^{p53-/-}$ cells after 48h siRNA treatment against SETDB1. GAPDH was used as a loading control. **B.** RPE1$^{p53-/-}$ and RPE1$^{p53-/-CHAMP1-/-}$ cells were treated with the SETDB1 inhibitor (Mithramycin A) for 24 hours, followed by a 20-minute EdU pulse. Cells were then treated with either 4 mM HU or DMSO for 2 hours. The number of H3K9me3-EdU PLA spots per nucleus was quantified. More than 50 cells were counted for each of three independent experiments. P values were calculated using one-way ANOVA, ****P<0.0001.

### Supplemental Figure 4

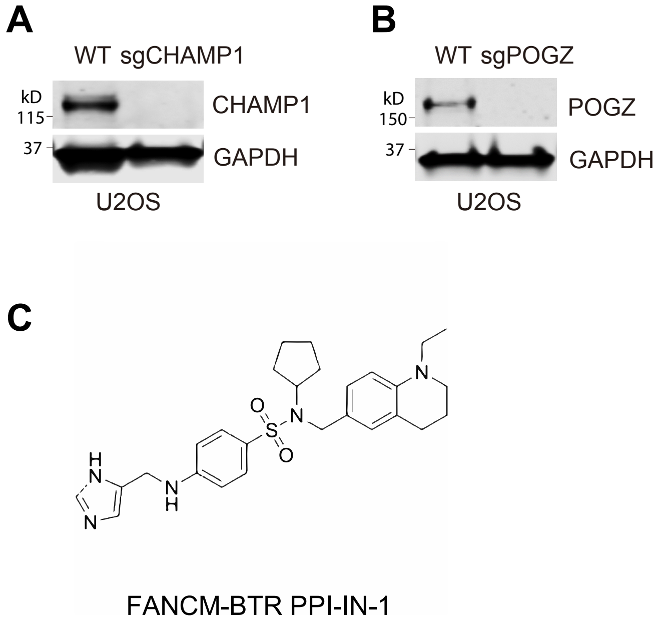


**Supplemental Figure 4, related to Fig. 4. A.** Immunoblot analysis of CHAMP1 expression in U2OS wild-type (WT) and CHAMP1 knockout cells. GAPDH serves as a loading control. **B.** Immunoblot analysis of POGZ expression in U2OS wild-type (WT) and POGZ knockout cells. GAPDH serves as a loading control. **C.** Chemical structure of FANCM inhibitor FANCM-BTR PPI-IN-1.

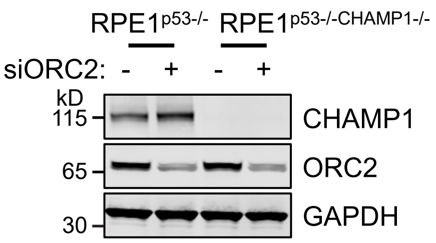


**Supplemental Figure 5, related to Fig. 5.** Immunoblotting against CHAMP1 and ORC2 in RPE1$^{p53-/-}$ and RPE1$^{p53-/-CHAMP1-/-}$ cells after 48h siRNA treatment against ORC2. GAPDH was used as a loading control.

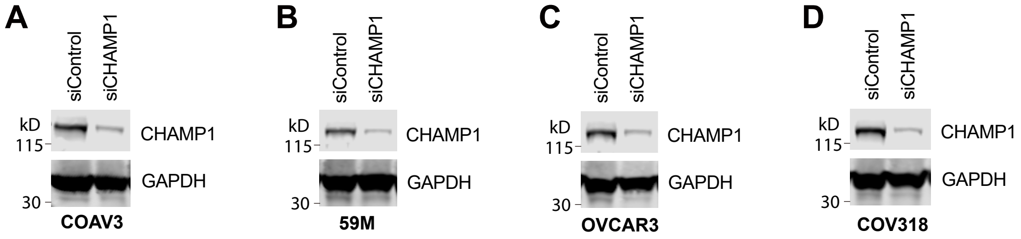


**Supplemental Figure 6, related to Fig. 6. Cancer cells with replication stress are highly dependent on CHAMP1. A-D.** Immunoblotting against CHAMP1 in COAV3 (**A**), 59M (**B**), OVCAR3 (**C**) and COV318 (**D**) cells after siRNA treatment against CHAMP1. GAPDH was used as a loading control.
